# Active control of arousal by a locus coeruleus GABAergic circuit

**DOI:** 10.1101/412338

**Authors:** Vincent Breton-Provencher, Mriganka Sur

**Affiliations:** Picower Institute for Learning and Memory Department of Brain and Cognitive Sciences Massachusetts Institute of Technology Cambridge, MA, 02139

## Abstract

Arousal and novelty responses linked to locus coeruleus noradrenergic (LC-NA) activity affect cognitive performance. However, the mechanisms that control modes of LC-NA activity remain unknown. Here, we reveal a local population of GABAergic neurons (LC-GABA) capable of modulating LC-NA activity and arousal. Monosynaptic retrograde virus tracing shows that inputs to LC-GABA and LC-NA neurons arise from similar regions, though a few regions provide differential inputs to one subtype over the other. Recordings in the LC demonstrate two modes of LC-GABA responses whereby spiking is either correlated or broadly anti-correlated with LC-NA responses, reflecting anatomically similar and functionally coincident inputs, or differential and non-coincident inputs, to LC-NA and LC-GABA neurons. Coincident inputs control the gain of phasic LC-NA mediated novelty responses, while non-coincident inputs, such as from the prefrontal cortex to LC, alter overall levels of LC-NA responses without affecting response gain. These findings demonstrate distinct modes by which an inhibitory LC circuit regulates the gain and tone of arousal in the brain.

## Introduction

Noradrenergic (NA) neurons located in the locus coeruleus (LC) send broad projections to a wide variety of brain regions, and their activity correlates with levels of arousal and cognitive performance^1-6^. During wakefulness, fluctuations in LC-NA arousal modify plasticity^7^, shift attention^8-10^, induce anxiety^11^, or affect discrimination and general sensory perception^1, 12-17^. Even though the effect of NA-mediated arousal on brain processing has become clearer in recent years, we still have a poor understanding of the mechanisms that regulate NA activity.

LC neurons receive inputs from a large number of brain regions^18^, which likely underlies the diverse contexts that drive LC-NA neuronal activity, including sensory stimuli and stressors^1-3, 7, 9, 11, 19-26^. In particular, LC neurons are thought to be novelty detectors, since NA release increases in response to novel sensory stimuli and with stimulus saliency^7, 19-22, 24-26^. The novelty response also changes with learning, suggesting that distinct mechanisms suppress or promote LC-NA mediated arousal^20-24^. Alongside these mechanisms regulating phasic LC responses, the modulation of tonic LC activity can also occur over longer time scales, such as during different levels of vigilance^3^, or during environmental changes^7^ or goal-directed behaviors^12^. Tight regulation of the global level of arousal has a key role in brain processing since unregulated arousal leads to hyperanxiety and detrimental performance^15, 27, 28^. One hypothesis to explain how LC responses are modulated is that local inhibition plays an active role in controlling LC-NA tonic activity and phasic responses. Consistent with this idea, previous *ex-vivo* reports using ultrastructural microscopy and slice recordings have shown that LC neurons receive direct GABAergic inputs, and it has been speculated that this inhibitory contribution originates from local GABAergic (LC-GABA) neurons^29-34^. Recordings across sleep-wake cycles have shown that GABA neurons in or near the LC are modulated during the sleep-wake cycle, similar to LC-NA neurons^35-37^. However, there has been no study of the location or function of LC-GABA neurons, the inputs they receive, or how they modulate LC-NA activity in the awake animal.

Here, we use a combination of anatomical, electrophysiological and optogenetic tools to identify the location, inputs and function of a local population of LC-GABA neurons in mice, and the mode by which they control LC-NA neurons. We find that the pattern of inputs to LC-GABA neurons allows two modes of inhibition of LC-NA activity: coincident inputs to LC-NA and LC-GABA neurons regulate the gain of phasic NA responses, whereas non-coincident inputs, such as preferential inputs from the prefrontal cortex (PFC) to LC-GABA neurons, regulate LC-NA tonic activity. Together, our findings identify a mechanism by which NA-mediated arousal is selectively modulated in the brain.

## Results

### Location of GABAergic neurons of the LC

Previous EM ultrastructural studies proposed the existence of GABAergic neurons surrounding LC^30, 33^. However, a clear map of the location of GABA neurons with respect to LC-NA neurons is lacking. To mark the precise location of LC-NA neurons, we injected dopamine-beta-hydroxylase (Dbh)-Cre mice with a Flex-tdTomato virus and examined coronal sections of the entire LC (**Fig. 1a**). We used immunohistochemical staining against GABA to localize LC-GABA neurons (**Fig. 1b**) and marked the location of LC-GABA with respect to LC-NA neurons in each mouse (**Fig. 1c**). Quantification of neuronal density in the LC region showed a greater density of LC-GABA neurons in the anterior and medial part of the LC (**Fig. 1d, e**). Overall, we observed that GABA neurons intermingle with and surround NA neurons in the LC.

**Figure 1.**
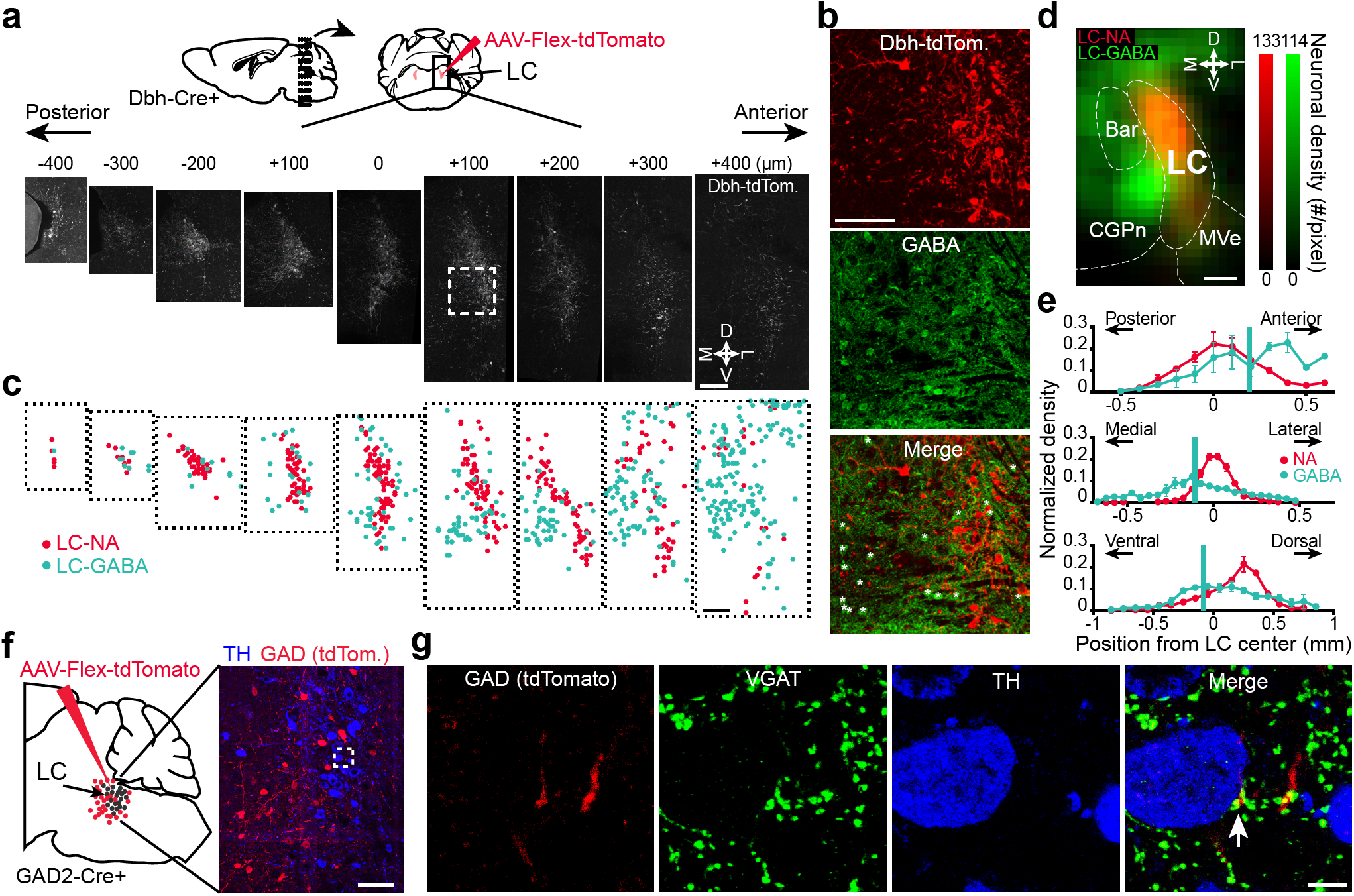
GABAergic neurons surround and contact LC noradrenergic neurons. **a.** Dbh-Cre mice were injected with AAV-Flex-tdTomato virus in the LC. Coronal sections around the LC were examined to reconstruct the complete structure of the LC-NA nucleus. Boxed area (at +100 um location) is shown magnified in (**b**). M: medial, D: dorsal, L: lateral; V: ventral. **b**. For each serial section, the location of GABA expressing somata (*) were revealed by immunohistochemistry. **c.** LC-NA and LC-GABA soma locations derived from (**a**) and (**b**). **d**. Map of LC-GABA and LC-NA neuronal density in a radius of 200 μm around the LC region. The density of each neuronal subtype was binned in 50 μm squares. The map is projected onto an antero-posterior axis. Bar: Barrington’s nucleus; MVe: medial vestibular nucleus; CGPn: central gray of the pons. **e**. Distribution of LC-GABA and LC-NA neurons in the 3 axes. Vertical green lines represent the center of the distribution for the GABA+ population. Data are presented as mean ± s.e.m. N = 3 mice. **f.** GAD2-Cre mice were injected with AAV-Flex-tdTomato virus to infect LC-GABA neurons. Coronal slices were stained for TH to reveal the location of LC-NA somata. Boxed area is shown magnified in (**g**). **g**. Example of an axon from a tdTomato expressing LC-GABA neuron apposed to a TH expressing soma in the LC (arrow). Staining against VGAT reveals the presence of presynaptic GABAergic terminals. Scale bars: (a), (c), and (d): 200 μm; (b) and (f): 100 μm; (g): 5 μm.

We next examined whether these local GABA neurons contact LC-NA neurons. We injected the LC of GAD2-Cre mice with a Flex-tdTomato virus to locate axonal processes from local GABA neurons, and marked LC-NA neurons by immunohistochemical staining against tyrosine hydroxylase (TH) (**Fig. 1f**). GABAergic processes entering the LC-NA region were apposed to TH expressing somas, and staining against the vesicular GABAergic transporter (VGAT) revealed the existence of GABAergic contacts (**Fig. 1g**). These results thus show that a dense population of GABAergic neurons is located in the LC region, and these neurons form putative synaptic contacts with LC-NA neurons.

### LC-GABA neurons reduce LC-NA-mediated arousal

To test if LC-GABA neurons functionally inhibit LC-NA neurons, and if they affect LC-mediated arousal, we first established the relationship between LC-NA activity and pupil size of awake head-restrained mice (**Fig. 2a, b**). We examined whether pupil size correlates with NA activity using two-photon imaging of axons filled with the genetically encoded calcium indicator GCaMP6s in the visual and prefrontal cortex of Dbh-Cre and TH-Cre mice (**Supplementary Fig. 1 a, b**). We found that NA activity correlated positively with pupil size and increased before pupil dilation events (**Supplementary Fig. 1 c-f**). We also confirmed these data with photo-identification of single-units recording in the LC of Dbh-Cre animals expressing Flox-ChR2 (**Supplementary Fig. 1 g, h**). As an additional indicator of arousal, we examined body movement along with pupil size and found a tight correlation between the two measures (**Supplementary Fig. 2**). This finding is consistent with previous studies which have shown that pupil size is correlated with cortical state and sensory processing^13-16^, and reflects LC neuronal activity^1, 5, 6^. We thus consider pupil size as a useful measure of NA-mediated arousal.

**Figure 2:**
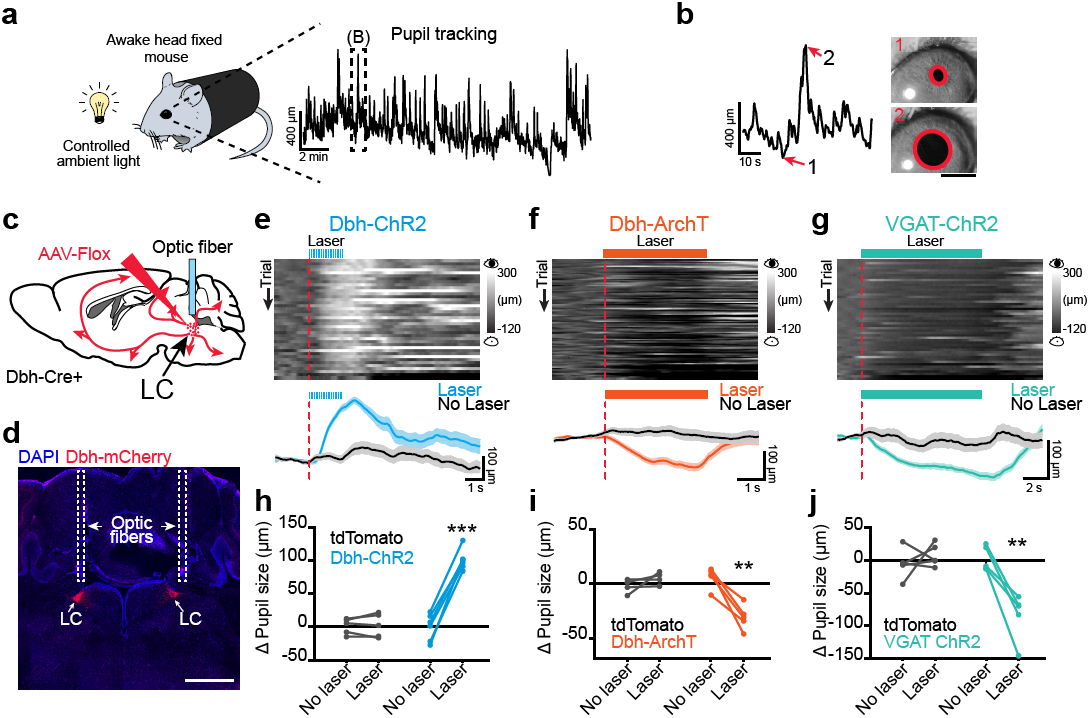
Activating LC-GABA neurons reduces LC-NA mediated pupil size. **a.** Pupil size was measured in awake head-fixed mice using a CMOS camera and infrared illumination. The trace on the right displays the pupil diameter for an example 20-minute session. Boxed area is expanded in (b). **b.** Example images of pupil tracking for constricted (1) and dilated epochs (2). Scale bar: 1 mm. **c.** Flox-ChR2-mCherry, Flox-ArchT-tdTomato, or Flex-tdTomato AAV constructs were injected bilaterally in the LC of Dbh-Cre mice. Optic fibers were implanted just over LC in both hemispheres and connected to laser light sources. For activating LC-GABA neurons, optic fibers were implanted in VGAT-ChR2-YFP mice. **d.** Coronal section of the LC showing ChR2-mCherry expression and optic fiber tracks. Scale bar: 1 mm. **e** – **g.** Effect of activating (Dbh-ChR2) or silencing (Dbh-ArchT) LC-NA neurons as well as activating LC-GABA neurons (VGAT-ChR2) on pupil size in example mice. Top panels – Temporal raster plots of pupil size aligned to optical activation onset (vertical red line). Bottom panels – session averages for trials with and without laser. Activation patterns were as noted in methods. Data are presented as mean ± s.e.m. **h** - **j.** Effect of activating or silencing LC-NA neurons as well as activating LC-GABA neurons on pupil size. Gray lines represent animals where only tdTomato was expressed, but similar optical activation patterns and intensities were used. Light activation patterns were the same as in (e-g). N = 6, 5 and 5 mice for ChR2, ArchT and VGAT-ChR2 conditions respectively. N = 5 mice for tdTomato controls. **: p < 0.01; ***: p < 0.001 using Student’s t-test.

To demonstrate causality between NA activity and pupil size, we injected a Flox-ChR2-mCherry virus bilaterally in the LC of Dbh-Cre mice (**Fig. 2c**), implanted optic fibers connected to a solid-state laser light source in the LC of both hemispheres (**Fig. 2c, d**), and applied pulsed blue light to mimic the observed increase in LC-NA activity preceding pupil dilation, as recorded in phototagged LC-NA units (**Supplementary Fig. 1g, h**). As hypothesized from NA axonal imaging, we observed a robust increase in pupil size after activation of ChR2-expressing NA neurons with blue light (**Fig. 2e, h**). Pupil dilation was dependent on baseline pupil size, since activating LC-NA neurons in periods of already increased arousal produced smaller increases in pupil size, thus explaining the observed trial to trial variability (**Supplementary Fig. 3a, b**). Even moderate activation of LC-NA neurons (pulse duration of 0.1 s at a frequency of 5 Hz) was sufficient to dilate the pupil, and this dilation increased with higher intensities of light activation (**Supplementary Fig. 3c, d**). We did not observe pupil dilation in control experiments using similar patterns of light activation in Dbh-Cre mice expressing only a fluorescent marker (Flex-tdTomato) (**Fig. 2h**).

**Figure 3:**
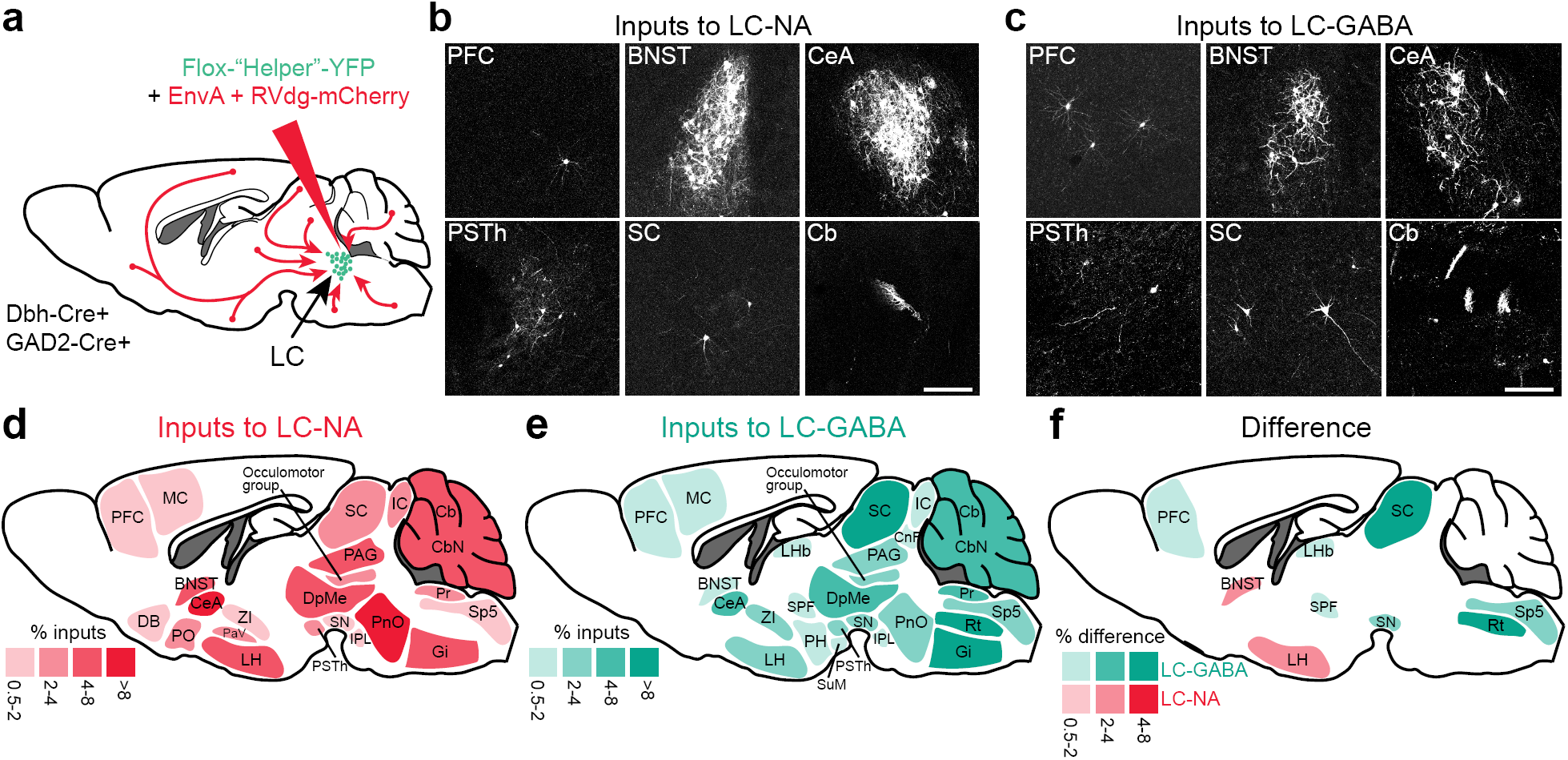
LC-NA and LC-GABA neurons receive inputs from similar as well as different sources. **a.** Schematic for targeting pseudo-rabies virus to LC-NA and LC-GABA subpopulation. **b,c.** Transynaptically labelled neurons in different brain regions following injection of targeted pseudo-rabies virus in LC of Dbh-Cre or GAD2-Cre mice. Scale bars: 200 μm. **d,e**. Map of brain regions providing the largest fraction of inputs to LC-NA and LC-GABA neurons. Regions providing less than 0.5% of total inputs are not displayed. N = 8 and 4 mice for LC-NA and LC-GABA respectively. **f.** Map of the difference in inputs to LC-NA and LC-GABA neurons. Only regions showing significant difference are displayed (p < 0.05 using paired t-test; see Table 1). BNST: bed nucleus of the stria terminalis; Cb: cerebellum; CbN: cerebellar nuclei; CeA: central amygdala; CnF: cuneiform nucleus; DB: diagonal band; DpMe: deep mesencephalic nucleus; Gi: gigantocellular nucleus; IC: inferior colliculus; IPL: interpeduncular nucleus; LH: lateral hypothalamus; LHb: lateral habenular nucleus; MC: motor cortex; PAG: periaqueductal gray; PaV: paraventricular nucleus; PFC: prefrontal cortex; PH: posterior hypothalamus; PnO: pontine nucleus; PO: preoptic nucleus; Pr: prepositus nucleus; PSTh: para-subthalamic nucleus; Rt: reticular nucleus; SC: superior colliculus; SN: substantia nigra; Sp5: spinal trigeminal tract; SuM: supramammillary nucleus; ZI: zona incerta.

Subsequently, we optically silenced LC-NA neurons to examine if their activity was necessary for pupil dilation. After injection of Flex-ArchT-mCherry virus in Dbh-Cre mice and implantation of optic fibers, we used a continuous pulse of green light to silence spontaneous LC-NA activity, and observed pupil constriction (**Fig. 2f, i**). While all tested mice showed pupil constriction when LC-NA neurons were silenced, this constriction was not observed in mice expressing the fluorophore alone (**Fig. 2i**). These results demonstrate that LC-NA activity is sufficient to alter pupil dilation and that a change in NA activity is reliably reflected in pupil size.

Our anatomical results suggested that local LC-GABA neurons are positioned to inhibit LC-NA neurons. We tested this hypothesis by implanting an optic fiber over the LC in VGAT-ChR2 mice, where all inhibitory neurons express ChR2^38^. Light activation produced a strong decrease in pupil size following focal stimulation, which was not observed in control animals (**Fig 2g, j**). Overall, these results demonstrate that local GABA neurons in the LC control the activity of NA neurons, and arousal level as reflected in pupil size.

### Inputs to LC-GABA and LC-NA neurons

To further understand the function of LC-GABA neurons with respect to LC-NA neurons, we targeted a modified rabies virus (RV) separately to each subpopulation to examine the anatomical sources of their presynaptic inputs^39^. We injected two Cre dependent (AAV) “helper” viruses to express the avian-specific retroviral receptor (TVA) and the rabies glycoprotein in GAD2-Cre or Dbh-Cre mice. Three weeks after the first injection, we injected the modified RV (rabies-deleted-glycoprotein-mCherry + EnvA) that only infected cells expressing TVA and spread only from cells expressing the glycoprotein (**Fig. 3a**). We waited an additional week and examined coronal sections of the entire brain from the olfactory bulb to 8 mm posterior from bregma. Both experiments showed neurons positive for mCherry in a wide range of locations (**Fig. 3b, c**; and **Supplementary Table 1**). We counted input neurons and assigned them to different brain regions by using a mouse brain reference atlas^40^. To visualize the projection patterns to LC-GABA and LC-NA neurons, we created a map of the relative contribution of each brain region to the two subpopulations (**Fig. 3d, e:** showing regions with > 0.5% total inputs). We excluded regions immediately surrounding the LC, as non-specific viral labeling of neurons occurred locally at the virus injection site. Brain regions contacting LC-NA neurons directly were similar to those reported previously, consisting of diverse structures related to sensory, cognitive, autonomic and motor functions^18^ (**Fig. 3d**). Nearly all of the approximately 50 regions providing major input to LC-NA neurons also projected to LC-GABA neurons (**Supplementary Table 1; Fig. 3e**). However, there were variations in the extent of projections; mapping the differential contribution of input regions to the two LC neuron subtypes showed 7 core regions projecting preferentially to LC-GABA versus 2 regions to LC-NA neurons (**Fig. 3f**). Thus, these findings demonstrate that the two neuronal subtypes in the LC receive largely similar inputs whereas a few regions provide preferential input to one neuronal subtype or another.

### Two types of arousal-related neurons in the LC

To understand how LC-NA and LC-GABA neurons integrate their inputs to influence arousal levels in the brain, we performed multi-unit recordings using a 16-channel silicon probe inserted in the LC of awake mice (**Fig. 4a**). The probe was coated with DiI crystals to verify the recording location, together with immunohistochemistry against TH to identify the LC, after each successful session (**Fig. 4b**). Alternatively, in a subset of recordings, we recorded from Dbh expressing neurons identified with optogenetics (phototagging). Dbh-Cre mice were injected with a Flox-Channelrhodopsin-2 (ChR2) virus. Four to six weeks after injection, we inserted a 16-channel silicon probe equipped with a 100 μm diameter optic fiber connected to a 473 nm laser source (**Fig. 4c**). Brief laser pulses (< 5 ms) were applied and units responding to laser pulses with a short delay (< 10 ms) were labeled as Dbh-ChR2 expressing (**Fig. 4d, e**). Sorting all recorded units by their spike waveform revealed two populations representing fast-spiking (FS) and regular spiking (RS) units based on their peak and valley width (**Fig. 4f**). The FS and RS unit populations represented 20.8% and 69.0% of the total units recorded respectively (the remaining units were not sortable using our criteria). The FS units showed higher levels of spontaneous activity and had shorter spike duration than RS units (**Fig. 4g, h, i**), matching previous recordings of verified GABAergic neurons of the pons^36^. Units responding to optical activation had similar waveforms and average firing rates as RS units (**Fig. 4g, h, i**). These data indicate that FS units correspond to putative LC-GABA neurons and RS units correspond to putative LC-NA neurons. The measurement of FS unit positions with respect to RS units showed a tendency toward FS units being located more ventral and anterior (**Fig. 4j**), though with considerable overlap, in a manner consistent with our immunohistochemical data. Overall, these results demonstrate that two relatively distinct types of units found within the LC likely represent GABAergic interneurons and NA neurons.

**Figure 4:**
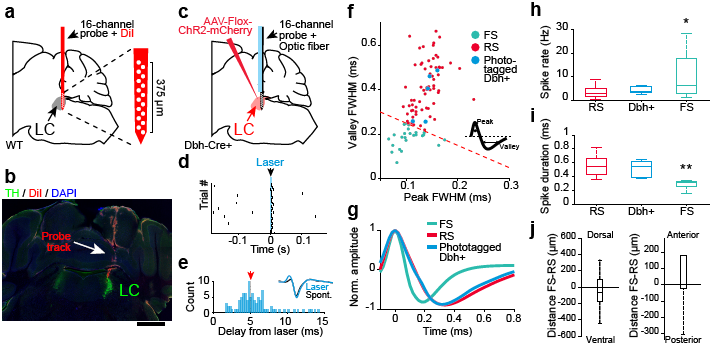
Extracellular spike recordings in LC reveal NA and putative GABA units. **a.** Single unit recordings were performed in the LC of awake mice by inserting a 16-channel silicone probe based on stereotactic coordinates. The probe was coated with DiI for later verification of recording sites. **b.** After each recording, the location of the silicone probe was verified with immunohistochemistry staining for TH to identify LC. Scale bar: 1 mm. **c.** In addition, LC-NA units were photo-tagged by inserting a 16-channel silicone probe equipped with a 100 μm optic fiber in the LC of Dbh-Cre mice previously injected with AAV-Flox-ChR2-mCherry virus. **d.** Spike raster for an example unit responding to 5 ms laser pulses. **e.** Distribution of delay from laser onset for the example unit in f. The red arrow represents the median of the distribution. Inset – Comparison of spontaneous versus laser-evoked spike waveforms. **f.** Classification of fast spiking (FS), regular spiking (RS) and phototagged Dbh+ units using peak and valley full width at half maximum values (see inset for definition). The red dashed line represents a threshold of 0.35 ms duration of each variable (n = 93 units from 16 mice). **g.** Average waveform for different types of units. **h, i**. Spontaneous spike rate and spike duration for RS, photo-tagged Dbh+ and FS units. *: p < 0.05 and **: p < 0.01 using ANOVA with Tukey post-hoc test. In (g-i), n = 69, 5 and 24 units from 16 mice for RS, photo-tagged and FS conditions respectively. **j**. Left - Measurement of the bias in the dorsoventral axis of FS with respect to RS units‥ Right – Measurement of the anterior-posterior bias of FS units with respect to RS units. No significant bias in the dorsoventral (p = 0.2638 using Student’s t-test, n = 24 FS units) or anterior-posterior axis (p = 0.0526 using Student’s t-test, n = 24 FS units) was found. Box plots indicate the median (center line), first quartiles (box edges) and minimum/maximum values (whiskers).

We next examined the relationship of RS and FS units to pupil size (**Fig. 5a**). As expected from the axonal calcium imaging data from LC-NA neurons (**Supplementary Fig. 1a-f**) and from responses of phototagged LC-NA units (**Supplementary Fig. 1g, h**), pupil dilation was associated with an increase in activity of RS and phototagged Dbh-ChR2+ units. In contrast, the activity of FS units was more heterogeneous, with individual units showing negative or positive correlation with pupil size (**Fig. 5a**). The Pearson correlation coefficient between firing rates of LC units and pupil size (**Fig. 5b**) showed that most of the RS units (87%) correlated positively with pupil size (**Fig. 5b, c**). This matched the correlation recorded with two-photon calcium imaging of LC-NA axons in the cortex (**Supplementary Fig. 1c**) and also the correlation of Dbh-ChR2+ units (where 5/5 units were positively correlated). On the other hand, FS units correlated both positively (FS+) and negatively (FS-) with pupil size (33% and 54% of FS units respectively; **Fig. 5c**). The activity of FS-neurons globally increased during periods of low pupil constriction while FS+ and RS neurons increased their activity during pupil dilation (**Fig. 5d**).

**Figure 5:**
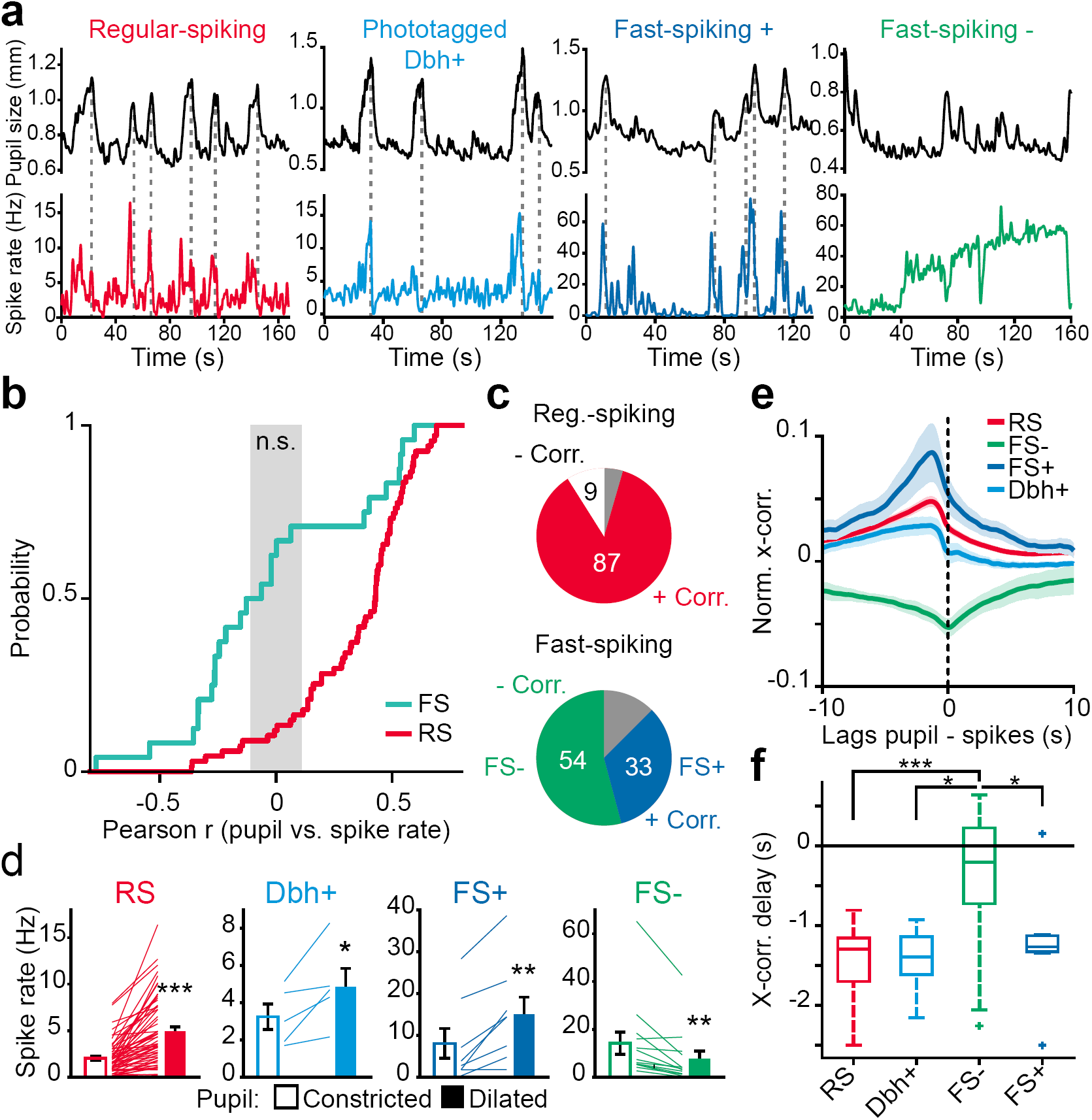
Putative LC-GABA units display two types of activity with respect to pupil size. **a**. Examples of simultaneous recordings of pupil size and RS, phototagged Dbh+ or FS LC units. Note the FS unit in the middle right panel is positively correlated while that in the far right panel is negatively correlated with pupil size. **b**. Cumulative probability distribution of the Pearson correlation coefficient of LC unit spike rate with pupil size for all FS and RS units. The gray area marks the region where Pearson correlation coefficients are non-significant. **c**. Percentage of RS and FS units that are positively or negatively correlated with pupil size. The gray portion indicates units not significantly correlated with pupil size. FS- and FS+ denote fast-spiking units with negative and positive pupil correlation respectively. Proportions were significantly different for RS versus FS group (χ2: 25.9284; p < 0.00001). n = 24 FS and 69 RS units respectively from 16 mice in (b) and (c). **d**. Average spike rate for different type of units during periods of constricted and dilated pupil, defined by values lower or higher than the 15th and 85th percentile of pupil size values respectively. Each line represents a single unit. *: p<0.01 **: p < 0.01; ***: p < 0.001 using paired Student’s t-test. **e**. Normalized cross-cor-relation of pupil size to LC firing activity for the classes of units sorted in (d). **f**. Delay between pupil dilation and LC unit activity as defined by the lag at maximum cross-correlation value for RS and FS+ units (or minimum for FS-units). **: p < 0.01 using ANOVA with Tukey post-hoc test. Box plots indicate the median (center line), first quartiles (box edges), minimum/maximum values (whiskers) and outliers (+). In (d-f) n = 58, 5, 13 and 8 for RS, Dbh+, FS- and FS+ units taken from 16 mice.

We examined more closely the effect of the different types of units on pupil dilation by calculating the cross correlation between spike rate and pupil size (**Fig. 5e**). The peak in cross-correlation for RS, phototagged Dbh+ and FS+ units overlapped, whereas the FS-units showed a broad trough in cross-correlation (**Fig. 5e, f)**. To get a better understanding of the timing of different LC units, we aligned their activity to dilation (and constriction) events (**Supplementary Fig. 4a**). RS and FS+ units showed a significant increase in their activity preceding dilation, whereas FS-units showed a significant reduction in activity that extended during dilation (**Supplementary Fig. 4b-c**). As a net effect, FS+ and RS activity correlated tightly during dilation and arousal, whereas FS-activity was broadly anti-correlated to the rest of LC activity. These two types of FS units did not differ in their spontaneous firing rate, their waveform duration or their recording location (**Supplementary Fig. 5**). Thus, FS+ and FS-units do not necessarily represent separate subpopulations of interneurons, but rather subpopulations that differ in the inputs they receive.

Overall, the two types of activity recorded from FS neurons are consistent with the findings of tracing experiments demonstrating similar as well as preferential inputs to LC-NA and LC-GABA neurons (**Fig. 3**), and they lead to specific hypotheses about their role. In one case, regions providing similar inputs may drive both LC-GABA and LC-NA neurons leading to correlated activity in these neurons, and inhibition could act as a feedforward gain control mechanism for arousal (similar to that seen in the cortex with thalamic inputs^41, 42^). In the other case, LC-GABA neurons that are broadly anti-correlated to arousal may be driven by a different input source than LC-NA neurons, allowing for independent positive and negative control of arousal.

### Coincident inputs to LC-GABA neurons provide gain control of LC-NA neurons

Our anatomical and electrophysiological results imply that a subset of LC-GABA neurons receives coincident inputs with LC-NA neurons. This suggests that feedforward inhibition is present at the level of LC and that local GABA neurons provide gain control to LC-NA neurons. We assessed this hypothesis by presenting auditory tone pips that varied randomly in frequency and amplitude. Tone pips activate LC neurons^19, 20, 24-26^, potentially by direct inputs from the inferior colliculus, the reticular formation of the pons, and the gigantocellular nucleus of the medulla, all these regions provide similar magnitude of input to LC-NA and LC-GABA neurons, as revealed by our anatomical tracer experiments (**Fig. 3d-f**). Recordings from GCaMP6s-expressing NA axons in the cortex confirmed that LC-NA neurons respond to tone pips (**Supplementary Fig. 6a, b**). Unit recordings of RS and FS neurons (**Fig. 6a, b**) showed that both responded to tone pips (**Fig. 6c**). The response delay from the onset of tone pip was 55.6 ± 3.1 ms and 54.0 ± 18.0 ms for the recorded population of RS and FS units respectively. These data suggest that auditory stimuli activate principally coincident inputs to LC-GABA and LC-NA neurons.

**Figure 6:**
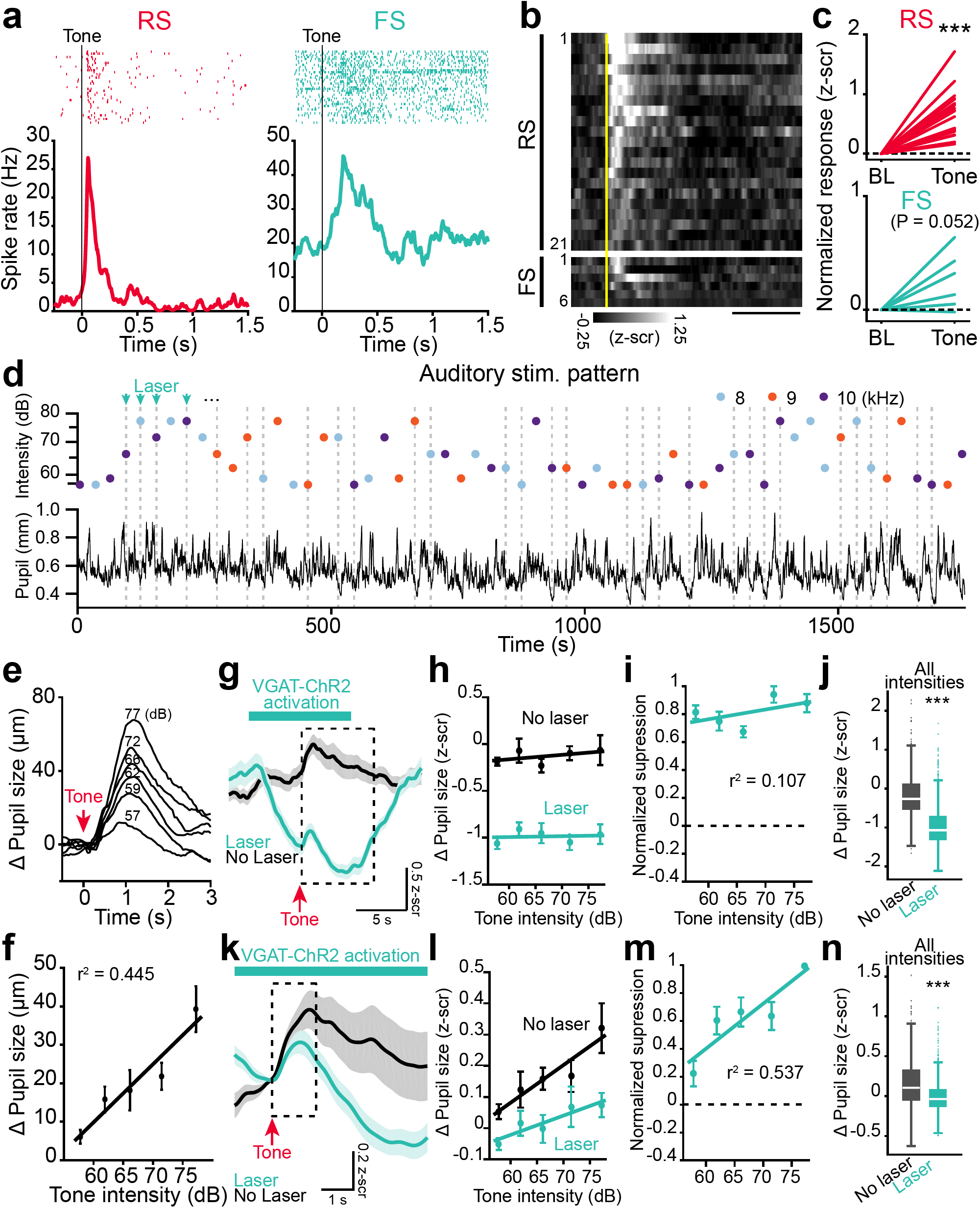
LC-GABA neurons control the gain of LC-NA mediated pupil response. **a.** Tone-pips (ranging from 8 to 10 kHz) at an intensity of 75 dB were presented while recording LC units previously classified as RS or FS based on their spontaneous activity waveforms. Left – regular spiking unit; Right – fast-spiking unit. Top panels – Spike raster plot aligned to tone onset for an example RS unit. Bottom panels – Session spike rate average. **b.** Raster plots of average responses to tone pips sorted between regular spiking and FS units. The vertical yellow line represents the tone onset. Scale bar: 0.5 s. **c.** Increase in firing rate following tone pips for all neurons in all categories. Note that population of RS units show significant increase to tone pip whereas FS units show almost significant increase (P values calculated using a Student’s paired t-test). In (b, c), n = 21 and 6 RS and FS units respectively. **d.** Top – Example of an auditory stimulus pattern used to test the effect of LC-GABA activation on LC-NA responses. Sound stimuli of 5 different intensities and 3 different frequencies were randomly presented at 30 sec intervals to VGAT-ChR2-YFP mice implanted with optic fibers in LC. Laser was activated during stimulus presentation on 50% of trials. Bottom – Corresponding pupil size measurement. **e.** Average of pupil size response to tones of different intensities for an example mouse. Each trace is an average of 32 to 70 trials depending on intensities. **f.** Change in pupil response amplitude with tone intensity (data for N=8 mice recorded in the normal condition, without laser activation). The line represents a linear fit (p < 0.0001 using one-way ANOVA). **g.** Average traces of trials where 72 dB tones were presented (red arrow) with and without laser activation of LC-GABA neurons for one mouse. The dashed box delineates the averaging window for calculating the tonic effect of VGAT-ChR2 activation (panels (h-j)). **h.** Pupil size at different tone intensities, with and without laser activation, for the example mouse in (g) using a 7-second averaging window after tone onset. **i.** Normalized suppression of tonic pupil response due to activation of LC-GABA neurons. Suppression was calculated by subtracting the response of laser on from the laser off trials and normalizing to maximum value. **j.** Comparison of laser to no laser trials for all trials regardless of tone intensity or frequency. **k**. Average traces for the same mouse and condition as in (g), showing the phasic post-tone-onset response. To compare response amplitudes, the traces were normalized to a baseline of 0.5 sec preceding the tone onset. The dashed box delineates the averaging window for calculating the phasic effect of VGAT-ChR2 activation (panels (l-n)). **l.** Pupil size at different tone intensities, with and without laser activation, for the same example mouse using a 1.5-sec averaging window after tone onset. **m**. Normalized suppression of phasic pupil response to tone intensity due to activation of LC-GABA neurons. **n.** Comparison of laser to no laser trials for all trials regardless of tone intensity or frequency. For panels (j) and (n) box plots indicate the median (center line), first quartiles (box edges), minimum/maximum values (whiskers) and outliers (+), n = 812 and 863 laser off and laser on trials respectively. For all other panels, data are displayed as mean ± s.e.m. N = 5 mice in (i), (j), (m), and (n). ***: p < 0.001 using unpaired Student’s t-test in (j) and (n).

Since tone pips evoked responses in both type of LC neurons, we used these stimuli to assess the possible modulation of LC-NA responses by LC-GABA neurons. We presented a random and sparse sequence of auditory stimuli of different frequencies and intensities to a group of mice, while recording pupil diameter to track the different NA responses (**Fig. 6d**). Simultaneous recording of pupil diameter and LC-NA activity showed that increase in pupil size followed the increase in LC-NA activity to auditory stimuli (**Supplementary Fig. 6b**). Silencing NA neurons prevented this pupil dilation response (**Supplementary Fig. 6c-d**), demonstrating that pupil size can be used to track NA activation with these stimuli. The pupil dilation response to auditory stimuli increased with the intensity of auditory tone independent of tone frequency (**Fig. 6e, f and Supplementary Fig. 7a-c**). Then, we used VGAT-ChR2 mice with fiber optics implanted in LC to examine the role of LC-GABA activity on LC-NA responses involving coincident inputs. The laser was turned on for 50% of the trials in randomized order (**Fig. 6d**). The trial averaged traces with or without laser activation demonstrated that activating LC-GABA neurons produced a constricting effect on pupil size, while mild pupil dilation was still detectable after the tone presentation (**Fig. 6g**). Similar to activating LC-GABA neurons during spontaneous fluctuations of arousal (**Fig. 2g,j**), we observed a uniform sustained (spanning several secs) reduction of pupil size across all sound intensities (**Fig. 6g-j**). For this experiment, we were interested in examining the effect of activating LC-GABA neurons on the transient response of LC-NA neurons to sensory stimuli. To remove the tonic effect of LC-GABA activation, we normalized the response for each tone to a 0.5 sec baseline period preceding the tone onset when the laser was already on, and averaged the pupil response to tone pips for a 1.5 sec period after baseline (**Fig. 6k**). Doing so, we observed that increasing the activity of LC-GABA neurons had a divisive effect on the tone intensity-pupil dilation relationship, as seen by the larger transient suppression of pupil size by laser activation at higher intensities and reduction of slope (or gain) of the pupil size change vs stimulus intensity line (**Fig. 6l**). This led to enhanced normalized suppression during presentation of tones of higher intensities independent of tone frequency (**Fig. 6m,n; Supplementary Fig. 7d**), similar to other examples of coincident feedforward inhibition^41, 42^. Altogether, these results demonstrate that activating LC-GABA neurons alone sets the baseline tone of pupil size, and that coincident activation regulates the gain of LC-NA mediated arousal.

**Figure 7:**
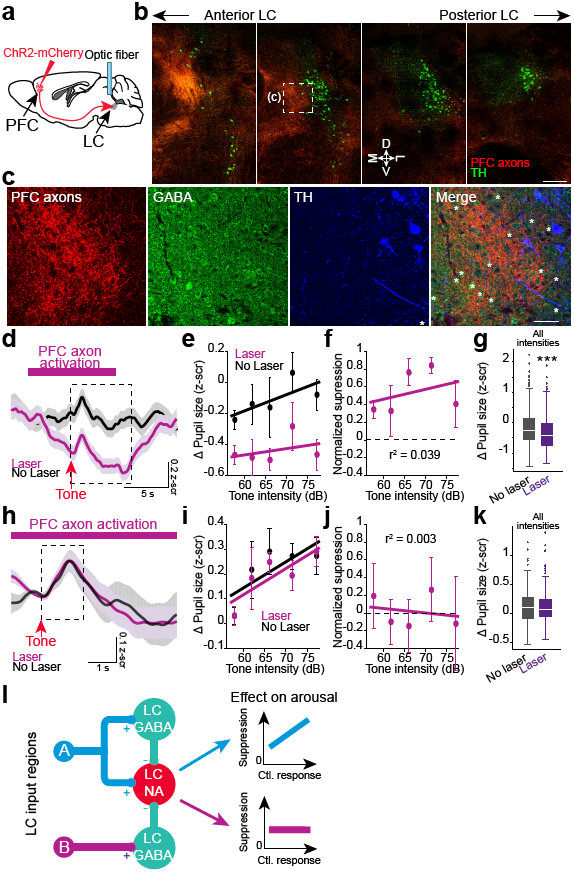
Projections from PFC to LC-GABA neurons control LC-NA mediated pupil tone. **a.** Methodology used to target PFC projections to LC. AAV-ChR2-mCherry was injected in the PFC and an optic fiber was implanted over the anterior part of LC. **b.** Confocal images of coronal sections from anterior (left) to posterior (right) LC showing PFC axons in LC and TH staining. Boxed area is shown in (c). M: medial, D: dorsal, L: lateral, and V: ventral directions. Scale bar: 200 μm. **c**. Confocal images of the region highlighted in b. Asterisks indicate GABA+ somata. Scale bar: 50 μm. **d.** Auditory tones (red arrow) were presented while activating PFC axons in LC. The traces represent the average of all trials for one mouse. The dashed box delineates the averaging window for calculating the tonic effect of VGAT-ChR2 activation (panels (e-g)). **e**. Pupil size at different tone intensities, with and without laser activation, for the example mouse in d using a 7-second averaging window after tone onset. **f.** Normalized suppression of tonic pupil response due to activation of PFC axons in LC. **g**. Comparison of laser to no laser trials for all trials regardless of tone intensity or frequency. **h**. Average traces for the same mouse and condition as in (d), showing the phasic post-tone onset response. To compare response amplitudes, the traces were normalized to a baseline of 0.5 second preceding the tone onset. The dashed box delineates the averaging window for calculating the phasic effect of PFC activation (panels (i-k)). **i**. Pupil size at different tone intensities, with and without laser activation, for the same example mouse using a 1.5-second averaging window after tone onset. **j.** Normalized suppression of phasic pupil response to tone intensity due to activation of LC-GABA neurons. **k**. Comparison of laser to no laser trials for all trials regardless of tone intensity or frequency. **l.** Proposed model of interaction between LC inputs and LC-GABA/LC-NA neurons. Excitatory coincident inputs from region A activate both LC-GABA and LC-NA neurons simultaneously. LC-GABA inhibition scales LC-NA activity divisively, thus controlling the gain of response. Non-coincident inputs from region B target LC-GABA neurons but not LC-NA neurons (as illustrated here), thus regulating overall NA tone without affecting the response gain. For panels (g) and (k) box plots indicate the median (center line), first quartiles (box edges), minimum/maximum values (whiskers) and outliers (+), n = 249 and 280 laser off and laser on trials respectively. For all other panels, data are displayed as mean ± s.e.m. N = 3 mice in (f), (g), (j) and (k). ***: p < 0.001 using Student’s t-test.

### Non-coincident inputs to LC-GABA neurons non-specifically suppress LC-NA neurons

The results from our retrograde labelling experiments suggested that along with similar, coincident, activation of LC-NA and LC-GABA neurons, some input regions also provide preferential, potentially non-coincident, drive to LC-GABA neurons. Moreover, our electrophysiological data showed that LC-GABA neurons displayed two types of activity, where the activity of FS+ neurons was correlated with LC-NA neurons while the activity of FS-neurons was broadly anti-correlated with LC-NA activity. We thus wished to examine if activation of non-coincident inputs to LC-GABA neurons would non-specifically suppress LC-NA activity, and hence regulate arousal tone.

We chose to target inputs from the prefrontal cortex (PFC), as our anatomical data supported the existence of a preferential drive to LC-GABA neurons originating from this region (**Fig. 3d-f**). We injected a virus expressing ChR2-mCherry in the orbitofrontal part of the prefrontal cortex and implanted an optic fiber above the LC to activate PFC axons (**Fig. 7a**). Coronal sections of the LC demonstrated the existence of PFC axons in the LC, confirming our results with monosynaptic retrograde labelling (**Fig. 7b, c**). Most of the PFC axons projected to the anterior and medial part of the LC, where the density of GABA neurons peaks (**Fig. 1e**; **Fig. 7c**). We activated PFC axons to the LC and indeed recorded significant sustained pupil constriction (lasting several secs) following laser activation (**Fig. 7d**). This constriction of pupil size was independent of tone intensity (**Fig. 7e-g**). Along with activating non-coincident inputs by PFC axonal activation, we activated coincident inputs to LC-NA and LC-GABA neurons with randomly varying auditory tone pips and recorded a transient increase in pupil dilation (**Fig. 7h**). However, analyzing the tone-induced pupil dilation (by normalizing to a baseline 0.5 sec before the tone presentation and by averaging the pupil response over 1.5 sec after tone pip onset) with and without laser activation revealed no effect of PFC activation on the gain of the transient pupil response (**Fig. 7i-k**). In contrast to the divisive effect of LC-GABA activation on pupil responses (**Fig. 6m**), the major effect of PFC activation was a non-specific reduction in pupil size (**Fig. 7g**), and hence a net decrease in arousal. Overall, these results demonstrate that non-coincident inputs such as from the PFC control the level or tone of LC-NA mediated arousal but not the gain, via their preferential targeting of LC-GABA neurons.

## Discussion

We demonstrate anatomically and functionally the existence of a local population of GABA neurons in the LC, which inhibits NA activity and thus controls arousal level in the brain, as reflected in pupil size. The pattern of LC afferents is mostly coincident to LC-NA and LC-GABA neurons (**Fig. 7l**), as shown by tracing experiments and correlated neuronal activity among the two subpopulations. This input pattern supports a role for local LC inhibition in controlling the gain of LC responses. On the other hand, some functionally distinct input regions exert stronger influence on one LC subpopulation over the other, suggesting that the general NA tone and accompanying arousal level can be set by these inputs (**Fig. 7l)**. An example of this second projection type is the PFC that projects preferentially to the LC-GABA sub-region and alters the tone of NA activity without altering its response gain.

Coincident and non-coincident inputs to LC-GABA neurons impact LC-NA modes of activity on many levels. By regulating the response gain of NA activity to novel stimuli, coincident inputs enable phasic LC activity to be maintained within a certain range, potentially restricting NA activity to an optimal level required for cognitive processing^15, 27, 28^. By preferentially targeting LC-GABA neurons, non-coincident inputs set thresholds for NA activation and enable modulation of tonic LC activity during different contexts. For example, PFC inputs involved in utility assessment can gate incoming sensory signals to convey the presence or absence of novelty via LC responses^1, 43^. Non-coincident drive to LC-GABA neurons can also impact attentional shifts by interfering with the ability of LC-NA neurons to respond to novel sensory stimuli. Indeed, recordings in the LC of behaving mice have shown adaptation of LC responses to novelty^7, 22, 23^and a switch of their response to different components of fear conditioning behavior^21^. Alterations in inhibition provided by the regions preferentially driving LC-GABA neurons can explain fluctuations in tonic LC activity observed within sessions in animals performing a visual discrimination task^12^, or during the sleep-wake cycle^3, 44^.

Our results show that LC-GABA neurons are an important source of inhibition for LC-NA neurons. However, we do not exclude other types of inhibitory mechanisms controlling LC activity. First, inhibition in the LC can originate from distal sources. Electrical stimulation of pontine and medulla nucleus such as the prepositus nucleus, pontine reticular nucleus and gigantocellular reticular nucleus yield significant reduction in the firing rate of LC neurons^45, 46^. However, these studies never confirmed if inhibition arose from direct inhibitory projection from these nuclei, or from disynaptic pathways such as from the preferential activation of local GABA neurons. In addition, retrograde labeling studies have shown that a significant GABAergic neuronal population from the posterior lateral hypothalamic area and the central amygdala projects directly to the LC region, but we do not know the function of these projections *in vivo*^*47*^. Second, inhibition in the LC can arise from NA mediated inhibition. A brief period of inhibition following the phasic response of LC neurons to sensory stimuli^19^ has been attributed, in part, to NA-mediated collateral inhibition, since blocking alpha-2 noradrenergic receptors moderately suppresses it^45, 48^. However, other mechanisms such as feedforward and feedback inhibition from neighboring LC-GABA neurons could also explained this post-activation inhibition. Finally, inhibition in the LC can originate from presynaptic release modulation. An example of this is κ-opiate receptors that colocalize with glutamate- and corticotropin releasing factor-positive axons in the LC^49^; activation of these receptors reduces the response of LC neurons to sensory stimuli^49^.

The effect of PFC activation on LC activity has remained controversial. Pharmacologically silencing the prefrontal cortex in rats increases LC activity^50^, while direct activation of the prefrontal cortex tends to also increase LC activity^50, 51^. It has been an open question whether the PFC sends direct descending excitatory or inhibitory projections to LC, and whether an inhibitory influence arises from an indirect pathway. Our monosynaptic tracing results (**Fig 3**) reconcile these studies, since both LC-NA and LC-GABA subpopulations receive direct inputs from the PFC, albeit with a significant preference for LC-GABA neurons. In line with the tracing experiments, we find in some animals a brief period of pupil dilation at the onset of laser activation, reflecting direct inputs to LC-NA neurons (**Supplementary Fig. 8a,b**). However, an extended period of activation of PFC axons in the LC shows a net pupil constriction effect, consistent with a greater influence on LC-GABA than LC-NA neurons.

Our results highlight the causal relationship between LC-NA activity and pupil dilation. The link between neuronal activity in the LC and pupil size had been previously established in different species^1, 5, 6^; however previous studies had not shown that specifically activating or silencing LC-NA neurons increases or decreases pupil size respectively. LC-NA neurons could control pupil dilation by direct inputs to parasympathetic and sympathetic preganglionic neurons^52, 53^. Importantly, our results demonstrate that LC-GABA neurons play an essential role in regulating LC-NA activity. In addition to maintaining the dynamic range of LC-NA responses via coincident feedforward inhibition, non-coincident inhibition driven by pathways such as from PFC to LC regulates the global tone of LC activity, and can potentially switch the mode of LC activity in a manner that is dependent on the internal state of the brain.

## Online Methods

Detailed Methods are described at the end of the document.

## Acknowledgements

We thank Ian Wickersham for his gift of the AAV helpers and dG-Rabies virus used for monosynaptic tracing. We thank Vincent Pham and Liadan Gunter for technical assistance on histology experiments. We thank Steven Flavell, Andrea Bari, Rafiq Huda, Grayson Sipe and other members of the Sur lab for helpful comments. This work was supported by NIH grant EY007023, FRQS postdoctoral training grant and NSERC postdoctoral fellowship program.

## Author contributions

V.B.-P. and M.S designed the experiments and wrote the manuscript. V.B.-P. carried out the experiments.

## Competing financial interests

The authors declare no competing financial interests.

## Methods

### Animals

All procedures were approved by the Massachusetts Institute of Technology’s Animal Care and Use Committee and conformed to NIH guidelines. Adult mice (> 2 month old) of either sex were used in this study. We used the following mouse lines for the specific expression of various viruses in noradrenergic, GABAergic, and cholinergic neurons: TH-Cre (B6.Cg-Tg(Th-cre)1Tmd/J, Jackson Laboratory), Dbh-Cre (B6.FVB(Cg)-Tg(Dbh-cre)KH212Gsat/Mmucd, MMRRC),, GAD2-Cre (Gad2tm2(cre)Zjh/J, Jackson Laboratory). Single-unit recordings of LC neurons were performed in C57Bl/6 wild-type mice. Optogenetic activation of LC GABAergic neurons (LC-GABA) was done on VGAT-YFP-ChR2 (B6.Cg-Tg(Slc32a1-COP4*H134R/EYFP)8Gfng/J, Jackson Laboratory).

### Stereotactic surgeries

All surgical procedures were performed under isoflurane anesthesia while maintaining body temperature at 37.5°C using an animal temperature controller (ATC2000, World Precision Instruments). After deep anesthesia was confirmed, mice were placed in a stereotaxic frame (51725D, Stoelting), scalp hairs were removed with hair-remover cream, the underlying skin was cleaned with 70% alcohol and betadine, and an incision was made in the scalp. The conjunctive tissue was removed by rubbing hydrogen peroxide on the skull. The skull was positioned such that the lambda and bregma marks were aligned on the anteroposterior and dorsoventral axes. A small hole was drilled through the skull at the location of interest. We used the following coordinates (according to bregma – in mm): LC: −5 to −5.2 anteroposterior, ± 0.9 mediolateral, and 2.9 to 3 dorsoventral; prefrontal cortex (PFC): 2.6 anteroposterior, ±1.3 mediolateral, and 2.0 dorsoventral. Viruses were delivered with a thin glass pipette at a rate of 100-200 nL per min by an infuser system (QSI 53311, Stoelting). The following viruses (titer: ∼10^-12^ virus molecules/mL) were injected for imaging and optogenetic experiments: Flex-GCaMP6s (AAV1.Syn.Flex.GCaMP6s.WPRE.SV40, U. Penn. Vector Core), Flox-ChR2-mCherry (AAV1.EF1.dflox.hChR2-(H134R).mCherry.WPRE.Hgh, U. Penn. Vector Core); Flex-ArchT-tdTomato (AAV2-CAG-Flex-ArchT-tdTomato, UNC Vector Core); Flex-tdTomato (AAV1-Flex-tdTomato, UNC Vector Core); and (AAV5/CamKIIalpha-hChR2-(H134R)-mCherry-WPRE-pA, UNC Vector Core). We delivered a volume of 400 to 500 nL per injection site in the LC of Dbh-Cre mice, and in the PFC of wild-type mice. We let mice recovered for 4-6 weeks after injection of virus for optimal opsin or calcium-indicator expression. For rabies monosynaptic tracing experiments, we injected rAAV1/SynP-DIO-sTpEpB helper virus in the LC of Dbh-Cre (500nL) or GAD2-Cre (100-200 nL) mice. 3 weeks after the first virus injection we injected the EnVA-RΔG.mCherry (400-500 nL) in the same location.

200 μm diameter optic fibers were implanted with the following procedures. After anesthesia and animal preparation, the scalp was removed, the skulled was cleared of conjunctive tissue and the neck muscles were retracted from the interparietal and occipital plates. After careful alignment of the skull, the optic fiber cannula was held by a stereotaxic manipulator and inserted at different locations (see virus injections for coordinates) slightly on top of (∼200 μm dorsoventral) the targeted region. Two-ferrule cannulas (TFC_200/245-0.37_4mm_TS2.0_FLT, Doric Lenses) were used for LC implantation. A single fiber optic cannulas (CFM12L05, Thorlabs) was implanted in LC for activation of LC PFC axons. For LC PFC axon activation, the cannula was implanted using the following coordinates to target the anterior and medial part of LC: −4.5 to −4.8 anteroposterior, ± 0.5 mediolateral, and 2.5 dorsoventral. The cannulas was attached to the skull with dental cement (Teets Denture Material, or C&B Metabond, Parkell). To avoid light reflection and absorption, dental cement was mixed with black ink pigment (Black Iron Oxide #18727, Schmincke). A custom-designed head plate^54^ was also implanted at the end of the surgery for head fixation. We performed fiber implantation 2 weeks after virus injection of opsin. For all surgeries, analgesic was given before and for 3 days following surgery.

To perform LC single unit recording in awake head-fixed mice, we implanted a head plate 1-2 weeks before recording. We prepared the animal using the same method as for fiber optic implantation. We used a custom design stereotactic arm to align the head plate parallel to the median and dorsal line of the skull during implantation. The head plate was attached to the skull using dental cement. The exposed skull was protected using rapid curing silicone elastomer (Kwik-Cast, WPI) topped with a fine layer of dental cement.

Two-photon calcium imaging was done through a cranial window. Following virus injection of GCaMP6s in the LC of TH-Cre or Dbh-Cre mice, we drilled a 3 mm circular window centered over the anterior part of V1 (∼ 3.5 mm posterior and ∼ 2 mm lateral to bregma) or the medial prefrontal cortex (∼2 mm anterior to bregma and centered on the midline). A 3 mm centered on a 5 mm coverslip (CS-5R and CS-3R, Warner Instruments), and glued together with UV adhesive (NOA 61 UV adhesive, Norland Products) was positioned over the craniotomy and attached to the skull using dental cement (C&B Metabond, Parkell). A custom made head plate was also attached to the skull for head fixation.

### Pupil and body movement monitoring

After fixing the mouse head using a previously implanted head plate, a high resolution CMOS Camera (DCC1545M, Thorlabs) camera equipped with a 1.0X telecentric lens (58-430, Edmund Optics) was pointed at either the left or right eye depending on the experimental set up. 780 nm infrared illumination was provided by a LED array light source (LIU780A, Thorlabs). Illumination was done at an angle of ∼60° for the corneal reflection spot to be cleared of pupil visualization. Video acquisition of eye images (240 x 184 pixels) was performed at 20 Hz by a custom-made MATLAB script. The ambient illumination was controlled by a 7” monitor (700YV, Xenarc Direct) placed 8 cm in front of the mouse and displayed a full field gray stimulus at an illuminance of ∼57 Lx (#403125, Extech Instruments). This level of ambient illumination was sufficient to keep pupil constricted within the space between the two eyelids. In all experiments, a master computer, controlling the visual and auditory stimuli, triggered pupil camera acquisition, as well as 2-photon imaging, optogenetic manipulations, or extracellular electrophysiology recordings via various data aquisition cards depending on the experiment (PCI-DIO24, Measurement Computing, NI USB-6259 or BNC-2110, National Instruments). Time-stamps of every pupil frame were saved for further alignment with imaging, electrophysiology or optogenetic experiments.

We segmented the images of a black pupil on a gray iris background by a sequence of image processing manipulations done with a custom-made MATLAB script. We adjusted the min and max pixel intensity of pupil images. We then normalized pixel values by the convolution of pupil frames by a 5*5 pixel kernel matrix of equal values. The images were binarized by a threshold value manually decided for each experiment. The binary images were filtered to extract the largest components located in the center of the image. The isolated binarized pupil image was then fitted with a least square fit of ellipse. From this fit we estimated the diameter of the pupil for each frame. The pupil segmentation was either done online, during the experiments, or offline on saved images of the pupil. Using this pipeline for pupil segmentation, very few frames had to be dropped due to poor fitting of the pupil. Most of the dropped frames were due to eye blinking or excessive micro-saccades and the diameter for these frames were estimated by interpolation.

To monitor body movement we used a 720p HD camera pointed at the mouse (LifeCam Cinema, Microsoft). The sampling rate used was 10 Hz. Movement data was synced with other experiments by displaying an indicator on the monitor. The movement metric was calculated by extracting the pixel data in time for regions around the nose, the neck, the paws and the ears of the animal. The mean difference in pixel value between each frame was calculated for each region and we normalized this value over a region of the image where no movements were expected. Periods of activity or quietness were isolated using a threshold value of 0.5 standard-deviation on the movement trace.

### Two-Photon imaging of LC-NA axons

4 weeks after virus injection, GCaMP6s+ noradrenergic axons were imaged in the cortex using a Prairie Ultima IV 2-photon microscopy system. Mice were head fixed and a light-shield was attached to their head plate, and the 25X/1.05 NA objective (XLPlan N, Olympus) was lowered on top of their cranial window. The 920 nm excitation of GCaMP6s was provided by a Ti:Sapphire tunable laser (Mai-Tai eHP, Spectra-Physics). Power at the objective ranged from 10 to 30 mW depending on GCaMP6s expression levels. After locating axons at 4X optical zoom, their activity was acquired at 5 frames per second for 10 minute blocks while simultaneously imaging the pupil using 8X optical zoom. Axons with significant signal to noise ratio were selected for analysis. We imaged 3 to 8 axons per mouse. The majority of our imaging field of view contained only one axon. After recording one field of view, we moved at least 1 mm away to find new axons. Care were taken to select axons from different branches, even though we cannot exclude the possibility that axons we considered as arising from different neurons were actually branches arising outside our imaging window. Using these criteria 31 axons were recorded from 6 mice.

After acquisition, time-lapse imaging sequences were corrected for x and y movement using template matching ImageJ plugins^55^. Multiple circular region of interests (3 to 7) were taken along each axon to extract the fluorescence intensity. The intensity in time for each region was then averaged together to minimize the noise in our signal. The dF/F signal was then calculated from this average by taking the mode of the signal as the reference value.

### Optogenetic modulation of LC and NB circuits

We used solid state laser illumination at 473 and 532 nm for activating ChR2 and ArchT respectively (MBL-III-473/1∼200 mW or MGL-III-532/1∼300 mW, Opto Engine LLC). A 200 μm / 0.39 NA patch cable (M72L02, Thorlabs) was connected to the laser output and to an intensity division cube (DMC_1×2i_VIS_FC, Doric Lenses) for bilateral LC. The patch cable (200 μm / 0.39 NA; M83L01 or M81L01, Thorlabs) was attached to the animal ferrule implant using corresponding ceramic mating sleeves. To block any light emitting from the interface between the patch cable and the implanted ferrule, a piece of black electrical tape was wrapped around the connection. The laser pulse duration, frequency and shape was controlled by a data acquisition system (Digidata 1440A, Molecular Devices) connected directly to the analog port of the laser power supply. For ChR2 activation of NA+, 10 ms pulses at a frequency of 3, 5, 10 or 30 Hz for 0.1, 0.5, 1, and 2 s were applied depending on the experiment. The peak power of every pulse was 4 to 5 mW at the laser output. For ChR2 activation in VGAT-ChR2-YFP mice, a sine wave at 25 Hz was used for various durations. A small ramp up (duration 0.5 sec) and ramp down (duration ∼ 5 sec) in power was applied on top of the sine wave. Low power light stimulation (peak power < 3 mW) was used for VGAT-ChR2-YFP mice to keep the spread of optical activation in the vicinity of the optic fiber end, thus increasing neuronal activity only in GABA neurons surrounding LC. For direct inactivation of LC-NA+ neurons with ArchT, 15 to 17 mW of power was applied for 4 seconds. The rebound effect of inactivation was prevented by ramping down the laser power at the end of each laser pulse for 1 sec. For activation of PFC axons in the LC, 10 ms pulses at a frequency of 20 Hz applied for 10 sec. The peak power for those experiments ranged from 12 to 18 mW. At the end of each experiments, the location of the optic fibers was verified with respect to neurons expressing the opsin in the brain. For VGAT-ChR2-YFP and LC-PFC experiments, location was also verified with respect to LC-NA by immunohistochemistry for TH.

### Single unit recordings

One or two days before the experiments, mice were head-fixed for 1 hr to habituate to head fixation. On the day of the experiments, the mice were anesthetized with isoflurane and the dental cement and silicone elastomer on the skull were removed. The mouse was placed on the stereotaxic frame and a 500 μm diameter craniotomy was performed on top of the recording site (from bregma: −4.9 to −5.4 mm anteroposterior and 0.6 to 1.1 mm mediolateral). The dura above the inferior colliculus was removed and the craniotomoy was protected with saline and a piece of gelfoam. The skull was covered again with silicone and the animal was allowed to recover for at least 2 to 3 hours for the anesthesia effect to washout completely. The awake animal was then headfixed and the silicone and gelfoam removed gently. 0.9% NaCl solution was used to keep the surface of the brain wet for the duration of the recordings.

After placing the animal in the recording set up, we submerged a reference silver wire in the NaCl solution on the skull surface. The position of the 16-channel silicone probe (A1×16-Poly2-5mm-50s-177-A16 or A1×16-5mm-50s-177-A16, NeuroNexus) was referenced on bregma and the surface of the brain and lowered slowly (1 min per mm) to 2.5 mm in the ventral axis using a motorized micromanipulator (MP – 285; Sutter Instrument Company). The extracellular signal was amplified using a 1x gain headstage (model E2a; Plexon) connected to a 50x preamp (PBX-247; Plexon) and digitized at 50 kHz. The signal was highpass filtered at 300Hz. LC units were identified from 2.5 to 3.5 mm from the surface of the brain. If the recordings were successful, the silicone probe was gently retracted and the recording tract was marked by re-entering the DiI coated probe (2 mg/mL – D3911, ThermoFisher Scientific) at the same location. The brain was harvested post-experiment and immunohistochemistry for confirming the probe location was performed. Spikes were isolated online with amplitude threshold using Plexon Recorder software, but re-sorted offline using Offline Sorter software based on their principal components, peak-valley and non-linear energy ratio.

For optical identification (photo-tagging) of LC-NA+ units, we used a solid-state blue laser (MBL-III-473/1∼200 mW, Opto Engine LLC) connected via a 105 μm / 0.22 NA patch cable (M61L01, Thorlabs) to an optic fiber glued to the 16 channel recording probes (A1×16-Poly2-5mm-50s-177-OA16LP, NeuroNexus). 2 to 5 millisecond-long light pulses at various light intensities (0.1 to 15 mW) were then repeatedly delivered in the tissue (Frequency: 1 Hz), and each channel was screened for light-evoked events. Spikes were considered as light responsive if they responded within 10 ms after light stimulus onset. We also kept only units responding to at least 75% of laser pulses, whose light evoked waveforms closely matched the spontaneous ones. During our recordings we noticed that high power illumination induced an electrical artifact in our recordings. This artifact was later removed by subtraction, since the artifact shape is constant from trial to trial.

### Sound stimuli

Tone pips were delivered using a single speaker (HK195, Harman/Kardon) located at a distance of 30 cm from the mouse. Sound stimuli were created and delivered using Psych-toolbox in MATLAB. Tone sequence was randomized in volume and frequency to avoid habituation of the response. The inter-stimulus interval was set at 30 seconds. The duration of each stimulus was fixed at 0.5 s. The speaker frequency range was calibrated using a USB calibrated measurement microphone (UMIK-1, Mini DSP) and the Room EQ Wizard software. The ambient noise in the room was estimated around 50 dB and the sound stimulus intensities were recorded by a sound level meter. To obtain the pupil response to different tone intensities and frequencies, an average of 63 ± 5 trials per tone intensity were recorded. To avoid long sessions and habituation, we limited each mouse to one 1-hour session per day. 5 ± 1 sessions were recorded per animal. For trials with laser activation of LC-GABA neurons in VGAT-ChR2 and PFC-LC implanted mice, the laser was turned on 5 s before the auditory tone onset and kept on for a total duration of 10 second (5 s baseline and 5 s after auditory stimulus). To calculate the effect of laser activation on NA tone, we used a 7-second long averaging window aligned to sound onset and compared the z-scored laser and non-laser trials. For measuring the modulation of pupil response created by laser activation, a baseline period of 0.5 second before tone onset was used to calculate the increase in pupil size to different tone intensities and the response over a 1.5-second-long averaging window aligned to sound onset was evaluated. Trials with high (2 times standard deviation) or low (1 time standard deviation) were excluded from the analysis.

### Histology

Under very deep anesthesia, mice were perfused transcardially with 0.9% NaCl followed by 4% PFA. The brains were harvested and post-fixed in 4% PFA at 4?°C overnight. In some experiments, brains were extracted without transcardial perfusion and only immersed in PFA overnight. Coronal sections (100-μm-thick) were cut using a vibratome (VT1200S, Leica) and were incubated overnight at 4 °C in PBS-T (0.2% Triton) + 3% bovine serum albumin and the following primary antibodies at 1:1000 dilution were used: chicken anti-tyrosine hydroxylase (TYH, Aves Labs), rabbit anti-VGAT (131002, Synaptic Systems), and rabbit anti-GABA (A2052, Sigma). We used the following secondary antibodies at a dilution of 1:500: Goat anti-Chicken 647 nm (A21449, ThermoFisher Scientific), Goat anti-Chicken 488 nm (A11039, ThermoFisher Scientific), and Goat anti-Rabbit 488 nm (A11034, ThermoFisher Scientific). Slices were mounted in Vectashield hard set mounting medium with DAPI (H-1500, Vector labs). The resulting immunofluorescence was imaged with a confocal system (TCS SP8, Leica) with 10X / 0.40 NA, 20X / 0.75 NA, or 63X / 1.40 NA objectives (Leica).

For the reconstruction of LC structures and surrounding areas, we collected the brain from Flex-tdTomato virus injected Dbh-Cre mice. We performed staining of GABA by immunohistochemistry on the slices covering a region of 1 to 1.5 mm on the anteroposterior axis that covered the full extent of the LC. The GABA+ and Dbh+ neurons of the LC were imaged by performing tiling reconstruction with a 20X objective zoomed 2X. We also took low magnification (10X) images of each slice for later registration across the different slices. The location of GABA neurons located within 200 μm of Dbh+ somas was then marked using ImageJ, and the whole LC was 3D reconstructed by exporting those values in a custom-made MATLAB program. All slices were aligned with respect to the center of mass of LC-Dbh neuron. To obtain the neuronal density of each LC-NA and LC-GABA neurons, we counted the number of somas in each bin (bin size: 50 or 100 μm) and normalized to the total number of somas counted.

For analysis of inputs to LC-NA versus LC-GABA populations, we collected the brain 1 week after injection of EnVA-RΔG.mCherry. 100 μm-thick coronal sections were produced, and one section every 200 μm was serially mounted on microscope slides. We analyzed sections from the middle of the olfactory bulb (+4.5 mm from bregma) to the end of the brain stem (−8mm posterior to bregma). Somata positive for mCherry were counted from all selected slices except for regions surrounding LC. The Paxinos and Franklin^40^ mouse atlas was used as a reference for identifying brain regions. We also verified our data against the Allen Mouse Brain Atlas^56^ and found similar results. Regions adjacent to LC were not considered for analysis due to non-specific expression of virus at the site of injection. The fraction of total inputs for each animal was obtained by dividing the number of mCherry+ soma for each brain region by the total number of somas counted. Animals with too few presynaptic neurons were not considered for analysis.

### Data processing

All data analysis, unless noted, was performed using custom-made MATLAB scripts. In single unit recordings, we found two classes of unit based on their waveform. For each unit, we normalized the average waveform from −1 to 1 and calculated the full width at half maximum of the peak and valley portions of the spike. Due to the challenge of recording single units in the LC of awake mice, the number of total units was too low (n = 115 total) to use clustering methods for extracting FS and RS units. We thus decided to use a fixed threshold value of 0.35 ms to classify unit as FS (< 0.35 ms) or RS (>0.35 ms) type. This empirical value matches what was previously reported in spike recordings from verified GABAergic neurons in the pons^36^. We calculated the spontaneous firing rate by averaging the number of spikes during a 10-minute period, where no sensory stimulus was presented. We excluded units with spontaneous activity above 10 Hz or below 1 Hz from the RS or FS spiking group respectively. The distance between FS and RS on a probe was obtained by averaging for an FS unit the distance with each RS on the vertical axis. A bias toward an FS unit to be more ventral from RS units was arbitrarily marked as negative. To track the distance of the FS and RS units in the antero-posterior axis, we imaged the entire coronal slice of the retrieved recording electrode location and referenced it on a brain atlas^40^to evaluate the posterior distance from bregma. Instantaneous spiking rate (*r(t)*) were obtained by using a kernel density estimation^57^ using the following equations:

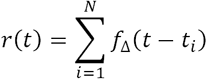

where *t*_*i*_ is the time if the *i*th spike and *N* is the total number of spikes. fΔ represent the following exponential kernel:

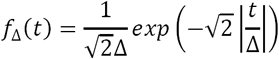

Different methods were employed to evaluate how neuronal or behavioral activity correlates with arousal. For measuring the general association between LC single unit activity and pupil size, we first obtained an estimation of the instantaneous spike rate sampled at 5 Hz. We then computed Pearson’s linear correlation coefficient between the pupil size and activity of LC. P-value for Pearson’s correlation were calculated using a Student’s t-distribution for a transformation of the correlation. A LC unit correlated significantly with pupil size if p < 0.05. The activity of different class of units during global pupil constriction or dilation was measured by averaging the spike rate during pupil values lower than the 25^th^ percentile (for constriction) or higher than the 75^th^ percentile (for dilation). To measure the timing of correlation with pupil, we computed the normalized cross-correlation. Pupil traces and neuronal or behavioral activity metrics were re-sampled (for extracellular LC recordings: 200 Hz, LC-NA Ca^2+^ axonal imaging: 5 Hz, and for facial movements video analyses: 10 Hz) and both normalized using z-score before computing cross-correlation. The delay between arousal and NA activity or movement was derived from the average lag value at maximum cross-correlation. To evaluate the increase in activity preceding pupil dilation, we band-pass filtered the pupil trace between 0.1 and 2 Hz using a second-order Butterworth filter. We then isolated pupil events by locating either the time point of minimas (dilation event) or maximas (constriction event) and extracted the pupil and spike rate traces for a window of −1 to 4 s around those time points. The amplitude of spike rate was evaluated from 0 to 0.5 after the onset of dilation or constriction. Alternatively, we also aligned NA activity (spike rate and calcium imaging) to maximas or minimas of the derivative of pupil size (variation of pupil diameter in time).

The effect of activating diverse populations of the LC on arousal was assessed by measuring the alteration in pupil size following light-activation or inactivation of neurons expressing ChR2 or ArchT. For each trial, we subtracted the baseline pupil diameter, evaluated for each trial on a period of 2 seconds before light onset. We then calculated the change in pupil size by averaging for a period of 1.5 to 8 s following the onset of the stimulus (Dbh-ChR2: 1.5 to 2 s depending on stimulus duration; Dbh-ArchT: 4 s and VGAT-ChR2: 8 s). Since there is a delay between increase or decrease in LC-NA activity and pupil dilation, we averaged the response of optical activation 0.7 sec after switching on the laser. The average variation in pupil size was then calculated for each animal. We used trials without laser as control to compensate for spontaneous changes in pupil size.

To evaluate how LC-NA and LC-GABA neurons respond to salient sensory stimuli, we recorded the response of LC-FS and RS units to tone pips. A unit was considered as auditory responsive if there was a significant increase of activity between a baseline period of 250 ms and the amplitude calculated over a 250-ms-period following the onset of tone pip. To evaluate the delay between neuronal response and stimulus onset, we calculated the average time for the response to reach a value above half the standard deviation. For two-photon imaging of LC-NA GCaMP6s+ axon, we compared the response with a 2-second baseline period preceding the auditory stimulus onset.

The effect of LC-GABA neurons on LC-NA mediated increase in arousal was assessed by recording the pupil diameter while animals were presented auditory stimuli of different intensities. We used VGAT-ChR2-YFP mice implanted with optic fibers in the LC on both hemispheres. LC-GABA neurons were optically activated by a 25Hz sine wave (max power ∼3 mW) on half the trials and the order of trials were randomized. Pupil response was calculated by subtracting a baseline pupil size value for each trial before the auditory stimulus (calculated over 0.5 s before tone onset). Since the effect of auditory stimulus on pupil dilation is usually delayed by a few ms, we calculated the average amplitude of pupil for a 1.5-second-window 0.5 second after the tone onset. This response was then averaged for each trial type to obtain the tone intensity – pupil size increase relationship. The suppression by LC-GABA neuron activation was defined as the difference between the response for trials with and without light activation. This difference was then normalized to the maximum value of suppression for each animal. The same procedure was used for the optical inactivation of LC-NA neurons with ArchT during sound stim. Optical silencing was performed for a total of 4 s (1s baseline and 3 sec post-stimulus) baseline and amplitude of pupil size was calculated for 1.4 and 1.6 s respectively surrounding the stimulus.

### Statistics

Throughout the paper we used paired and unpaired Student’s two-sided t-test for evaluating P values of experiments with two conditions. P values for experiments with multiple conditions were computed using ANOVA with Tukey post-hoc test. Significance level were marked as *: p < 0.05; **: p < 0.01; and ***: p < 0.001. P-value for Pearson’s correlation were calculated using a Student’s t-distribution for a transformation of the correlation. LC unit correlated significantly with pupil size if p < 0.05. Proportions of cells positively or negatively correlated with pupil for RS and FS units were tested for significance using a χ^2^ test. The activity of different class of units during global pupil constriction or dilation was measured by averaging the spike rate during pupil values lower than the 25^th^ percentile (for constriction) or higher than the 75^th^ percentile (for dilation). For all experiments, we used sample sizes to provide at least 80% power to detect an effect. Data is available upon request from the authors.

